# An improved protocol for high-efficiency and cost-effective CRISPR/Cas9-mediated knock-ins in *C. elegans*

**DOI:** 10.64898/2026.09.04.749330

**Authors:** Atal Vats, Arunima Sen, Ananya Bandyopadhyay, Murali Sudhanand, Marlyn Xavier Mascarenhas, Abhishek Bhattacharya

## Abstract

CRISPR/Cas9-mediated homology-directed precise genome editing using long single-stranded DNA (ssDNA) donors has expanded the possibilities for generating defined genetic modifications. However, the required ssDNA donor preparation can be technically demanding and often requires extensive locus-specific sequence design. Here, we examined parameters influencing ssDNA donor-mediated genome editing and developed approaches to simplify donor preparation using λ-exonuclease-mediated ssDNA generation. By evaluating donor designs across multiple genomic loci, we found that efficient genome editing can be achieved with relatively short homology regions for a range of insertion sizes. These findings provide a basis for simplifying ssDNA donor preparation and the overall gene-editing pipeline, potentially facilitating its application across species.

**Key features:**

1. This protocol offers cost-effective, alternative ssDNA donor preparation strategies, overcoming the requirement of expensive locus-specific, long phosphorylated primers.
2. Efficient genome editing can be achieved with relatively short homology regions for a range of insertion sizes.
3. This protocol provides a simplified CRISPR/Cas9-mediated knock-in pipeline suitable for editing a large number of loci.

**This protocol is used in:**

1. Rosenkranz, N., et al., *In situ structure of a gap junction-stomatin complex*. Sci Adv, 2025.
2. Vats, A., et al., *The combinatorial innexin code of heterochannel electrical synapses governs synaptic function and is maintained by distinct cellular mechanisms*. Proc Natl Acad Sci U S A, 2026.

## Background

CRISPR/Cas9-mediated homologous repair has become the method of choice for generating precise genetic modifications across species. Several studies have described diverse methods for the efficient introduction of both small (100 bp) and large (>1 kb) sequences into the genome, each offering specific advantages[1-8]. Among them, the introduction of sequence features (>150 bp) using single-stranded DNA (ssDNA) donor templates or hybrid dsDNA donor templates containing single-stranded homologous regions at the termini offers enhanced efficiency of successful editing events compared to traditional dsDNA donor templates [1, 4, 8]. Specifically, generation of long ssDNA donors (>150 bp) with homologous repair sequences in large quantities by selective digestion of one strand using λ-exonuclease has enabled high-efficiency genome editing in *Caenorhabditis elegans* [1]. This method first requires generation of a hemi-phosphorylated dsDNA donor by PCR amplification using one long, locus-specific, 5′-phosphorylated primer and one long conventional non-phosphorylated reverse primer, followed by preferential digestion of the phosphorylated strand by λ-exonuclease[1]. However, the requirement of these expensive locus-specific, 5′-phosphorylated primers for donor preparation increases both the cost and preparation time associated with each new genomic target. We therefore sought to simplify this workflow by developing alternative strategies to generate long ssDNA donors in large quantities, without requiring long, locus-specific 5′-phosphorylated primers. Secondly, we also sought to optimize the CRISPR reaction and microinjection protocol using these ssDNA donors to further reduce the cost and overall time required to obtain the edited progeny.

The use of long homology regions in the λ-exonuclease-mediated ssDNA generation approach presents a particular practical limitation wherein each new target requires synthesis of a new, 5′-phosphorylated primer containing the locus-specific 100 – 170 bp long homology sequence. However, previous independent studies suggest that homologous recombination-mediated knock-ins using significantly shorter homology regions have also been successful [2, 4]. Hence, we sought to develop ssDNA donor-generation strategies that incorporate locus-specific homology sequences without requiring long phosphorylated primers. We developed two alternative approaches to generate hemi-phosphorylated dsDNA substrates for λ-exonuclease-mediated digestion. In the first strategy, we introduced locus-specific, shorter 30 – 70 bp homology sequences on both sides of the desired insertion sequence using regular non-phosphorylated primers. This step also introduces unique 20 bp ‘adaptor sequences’ on both sides of the homology arms (Figure 1A). The resulting product was then used as the template to generate hemi-phosphorylated donor templates in sufficient quantity (> 8 µg) by a second PCR using one universal 5′-phosphorylated primer, complementary to the 5′ adaptor sequence, and another non-phosphorylated reverse primer complementary to the 3′ adaptor sequence. This design separates the locus-specific homology arm introduction step from the phosphorylation step, allowing the same phosphorylated primer to be used across different genomic targets.

**Figure 1.**
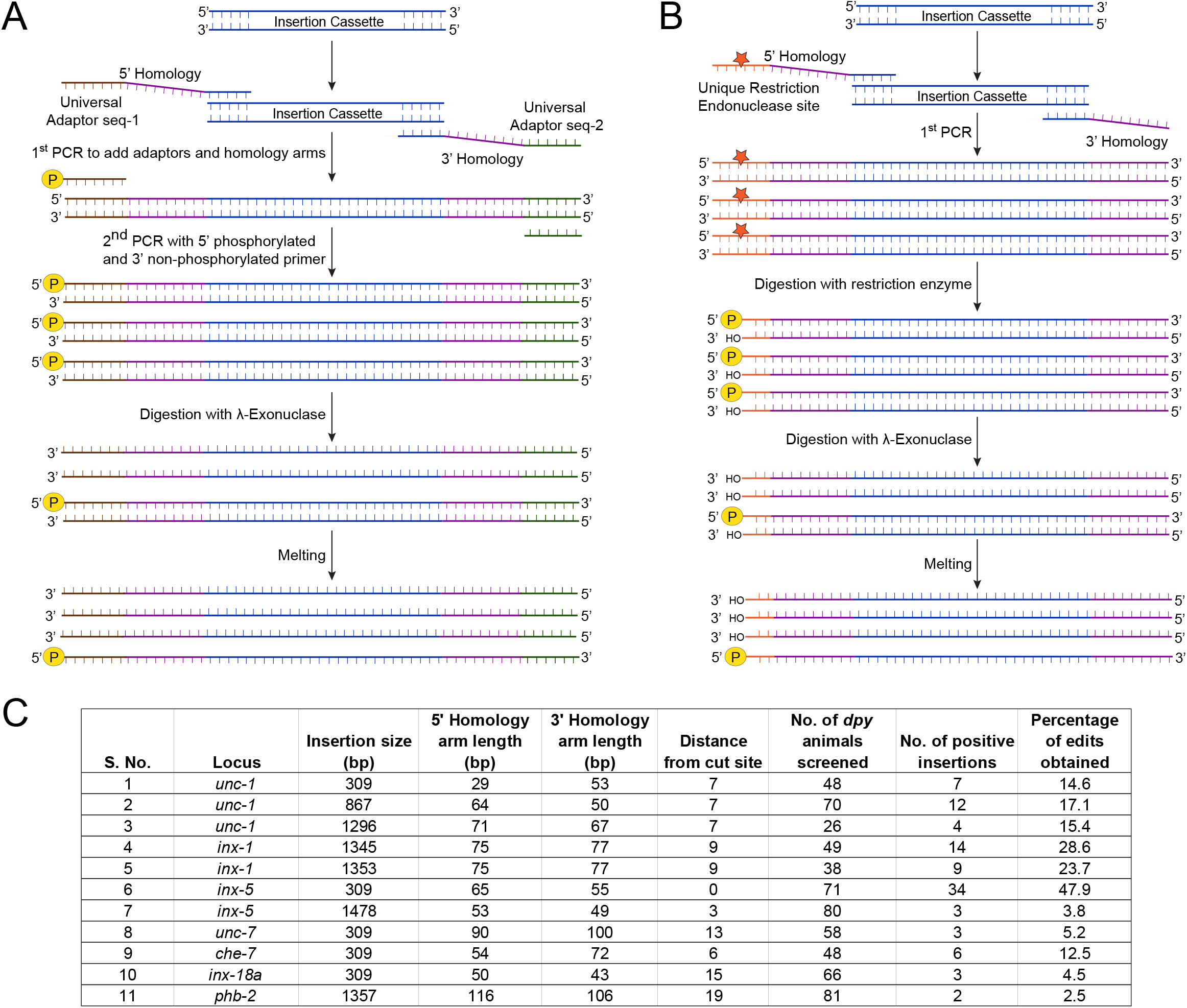
Alternative strategies for simplified, cost-effective ssDNA-donor-mediated homologous genome-editing. (A) Schematic of donor-generation Method 1. Gene-specific homology arms are fused to universal adaptor sequences flanking the insertion cassette in a first PCR. A second PCR uses a 5′-phosphorylated forward primer and a non-phosphorylated reverse primer to generate double-stranded product with one phosphorylated strand. λ-exonuclease selectively degrades the 5′-phosphorylated strand. Residual dsDNA is melted to yield enriched ssDNA donor. (B) Schematic of donor-generation Method 2. A unique restriction endonuclease site is incorporated adjacent to the 5′ homology arm in the first PCR product. Restriction digestion exposes a 5′-phosphorylated end, which is then selectively degraded by λ-exonuclease, followed by melting of residual dsDNA to generate enriched ssDNA donor. (C) Summary of 11 independent knock-in experiments across 7 loci (unc-1, inx-1, inx-5, unc-7, che-7, inx-18a, phb-2), listing insertion size, 5′ and 3′ homology-based repair arm lengths, distances of the insertion sites from the Cas9 cut sites, number of animals screened (*dpy-10* co-CRISPR-marked F1s), number of positive insertions, and percentage editing efficiency as validated by PCR genotyping.

In the second alternative strategy, we exploited the property of restriction endonucleases to generate a 5′ phosphorylated and 3′ hydroxyl end at the cut site to obtain hemi-phosphorylated donor templates. We incorporated a unique, blunt-end restriction endonuclease recognition sequence within one of the adaptor sequences introduced in the first PCR reaction (Figure 1B). Restriction digestion of the first PCR product with the corresponding endonuclease resulted in a hemi-phosphorylated double-stranded donor DNA containing a locus-specific homology sequence. The resulting hemi-phosphorylated dsDNA substrates were treated with λ-exonuclease to generate ssDNA donors. Together, these complementary strategies eliminated the need for a long, locus-specific 5′-phosphorylated primer, thus reducing material costs and procurement time.

λ-exonuclease treatment does not completely digest the dsDNA substrate and may leave residual dsDNA [1]. To further enrich the desired ssDNA donor population, we subjected the λ-exonuclease-treated DNA to an additional dsDNA melting step, as previously described [4]. Moreover, to facilitate the identification of genome-edited progeny from the injected animals, we also incorporated *dpy-10* co-CRISPR screening into our workflow [6].

Detailed procedures for donor preparation, λ-exonuclease treatment, injection-mix preparation, injection, and screening for genome-edited animals are provided below. Altogether, our findings identified a relatively simpler, cost-effective strategy for generating CRISPR/Cas9-mediated homology-directed knock-ins (Figure 3).

**Figure 2.**
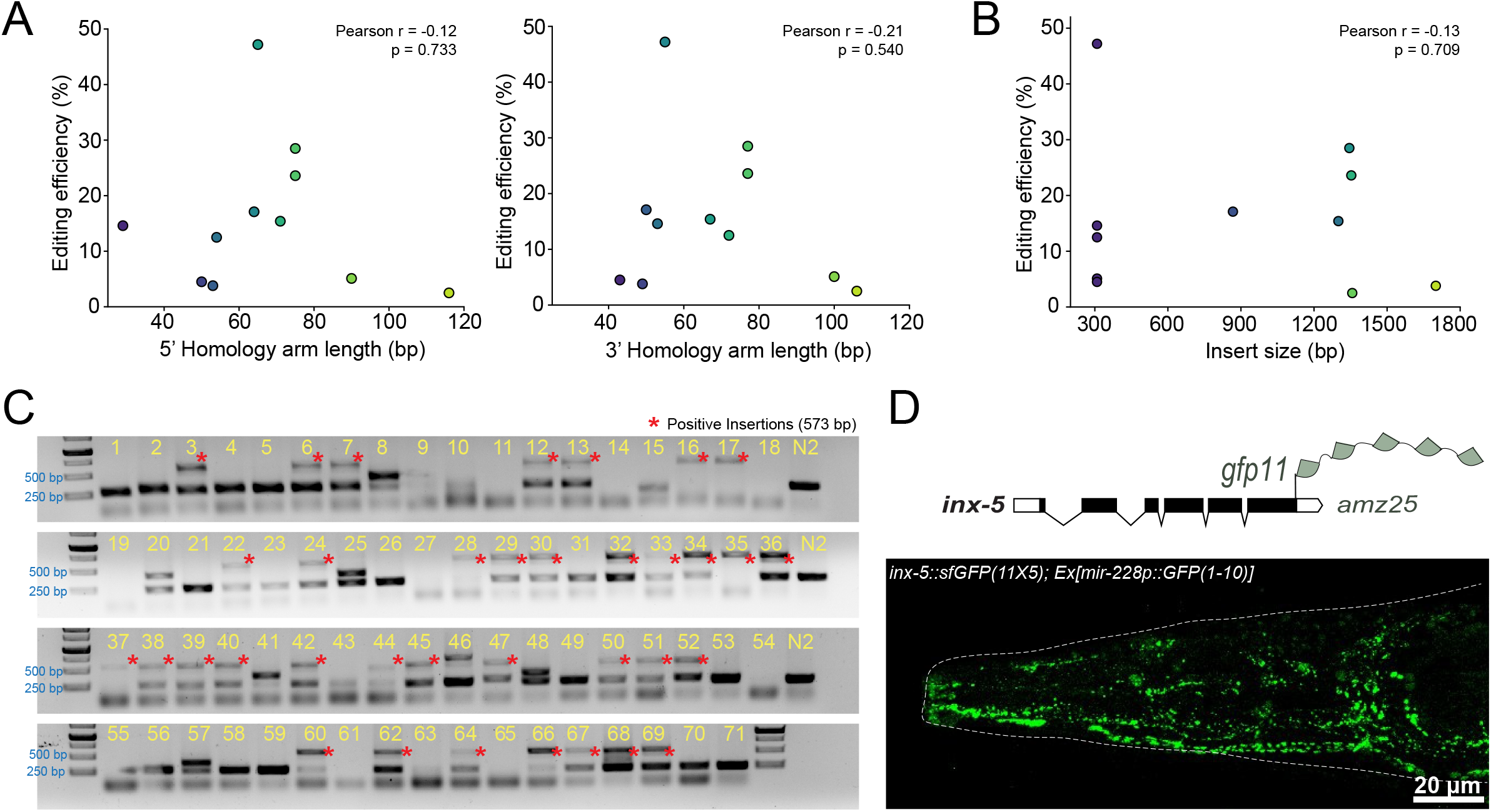
Comparison of editing efficiency across donor designs and validation of *inx-5* knock-in allele. (A) Genome editing efficiency (%) plotted against 5′ (left) and 3′ (right) homology arm length (bp) for the 11 insertions. Pearson correlation coefficients and p-values are shown; neither homology arm length showed a significant correlation with editing efficiency. (B) Genome editing efficiency (%) plotted against insertion size (bp) for the same 11 insertions. No significant correlation was observed. Pearson correlation coefficients and p-values are shown. (C) Representative agarose gels for PCR genotyping of the *inx-5* locus knock-in to generate *inx-5::sfGFP(11×5)*. Singled F1s (numbered 1–71) were genotyped. Asterisks (*) mark positive insertion band; N2 = wild-type control. Arrowheads mark the PCR bands corresponding to edited genome. (D) Top: Schematic of the *inx-5* locus showing C-terminal insertion of the gfp11 (split-GFP β-strand 11) tag, generating the allele *inx-5(amz25)*. Bottom: representative confocal image of an *inx-5::sfGFP(11×5)* knock-in animal expressing GFP1-10 under the pan-glial *mir-228* cis-regulatory element, showing reconstituted GFP fluorescence confirming correct in-frame insertion. Scale bar = 20 µm.

**Figure 3.**
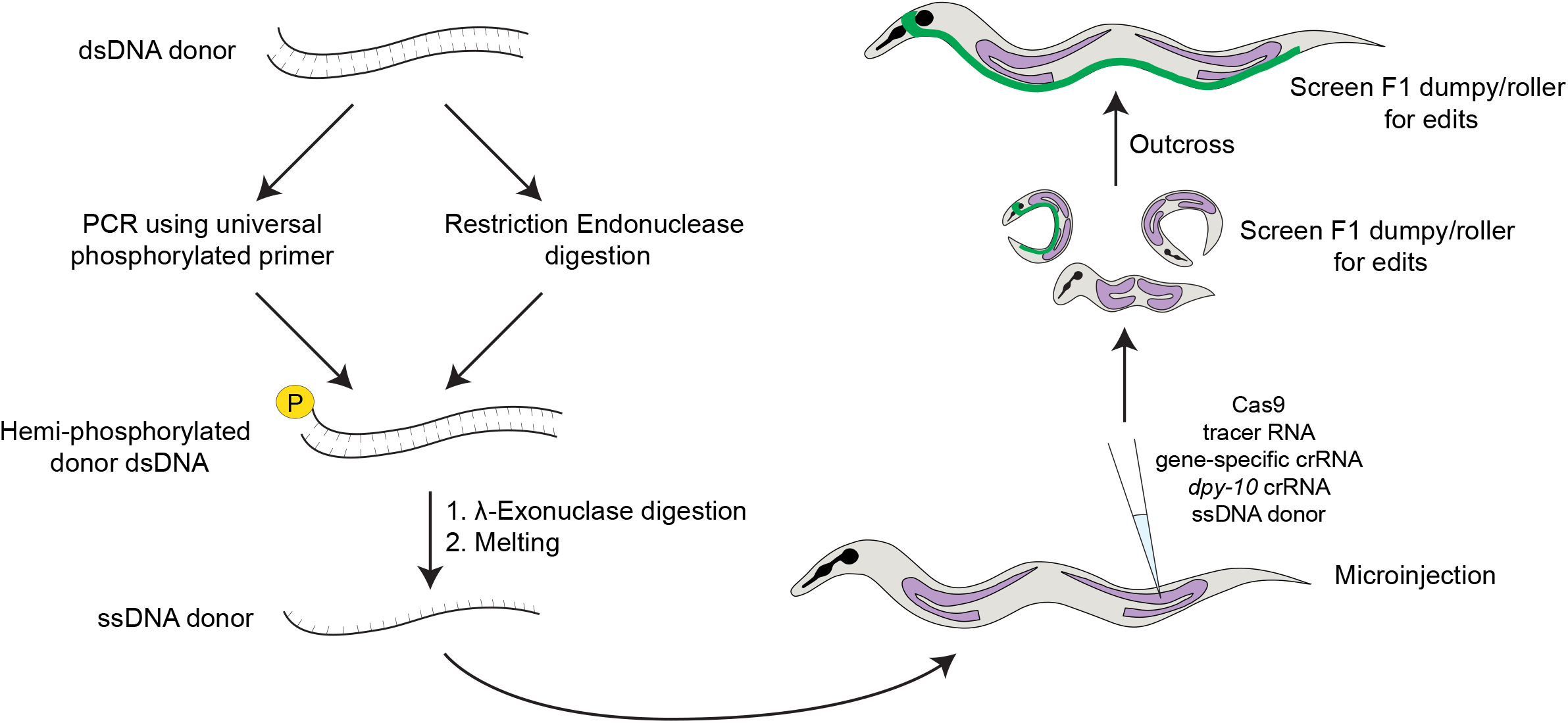
Graphical representation of the gene-editing protocol Schematics showing different steps of the homology-directed gene-editing protocol in *C. elegans*, with particular emphasis on the modified steps.

## Materials and reagents

### Biological materials

1. *Escherichia coli* OP50 (bacterial food source for *C. elegans* maintenance)
2. *Caenorhabditis elegans* strains generated in this study, their genotypes and corresponding crRNA sequences are listed in Table 1.

**Table 1.**
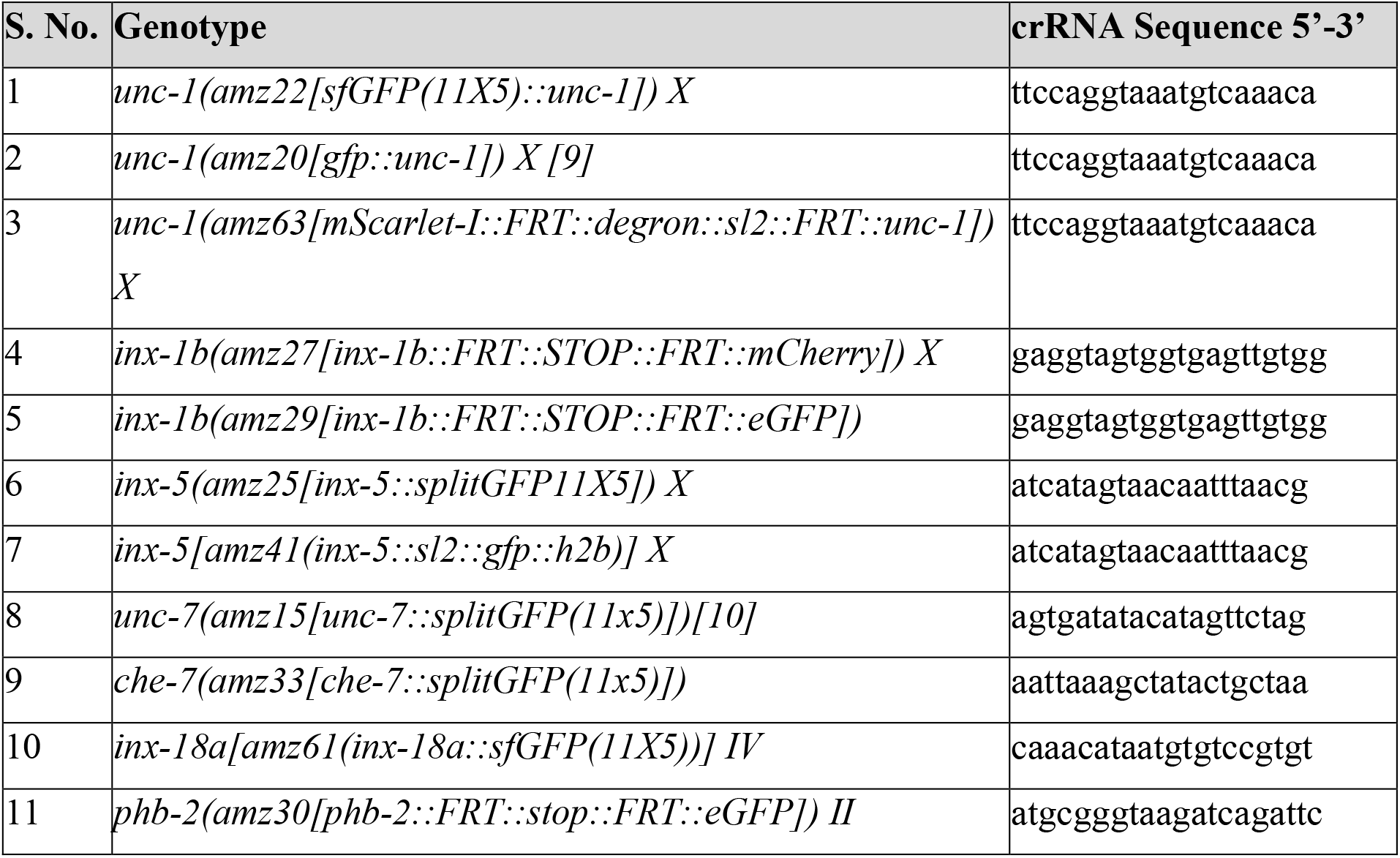
Strains and alleles generated in this study, with corresponding crRNA sequences.

### Reagents

1. Cas9 protein (IDT, catalog number: 1081059)
2. tracrRNA (IDT, catalog number: 1072533)
3. Gene-specific crRNA (IDT; custom synthesized per locus; sequences listed in Table 1)
4. *dpy-10* crRNA (IDT; sequence: 5′-gctaccataggcaccacgag-3′)
5. *dpy-10* ssODN repair template (IDT)
6. 10X λ-exonuclease reaction buffer (NEB, supplied with M0262)
7. λ-exonuclease (NEB, catalog number: M0262)
8. EDTA, 0.5 M (used to stop the λ-exonuclease reaction)
9. Nuclease-free H_2_O
10. Levamisole, 5 mM (used to immobilize animals for imaging)

### Laboratory supplies

1. QIAquick PCR Purification Kit (QIAGEN, catalog number: 28104)
2. Spin/purification columns for ssDNA clean-up (Any purification kit with a binding capacity >5 µg was effective)
3. Microinjection needles
4. Agarose pads (for immobilizing animals during imaging)

### Equipment

1. Thermal cycler (for PCR amplification and the DNA melting cycle)
2. Microinjection setup/microinjector for *C. elegans* microinjection
3. Microcentrifuge (capable of 15,000 rpm)
4. Incubators (20 °C, 25 °C, and 37 °C)

### Software and datasets

1. ImageJ Fiji (used to process and adjust fluorescence images for display)

### Procedure

#### A. *C. elegans* maintenance and strains

*C. elegans* strains were maintained on nematode growth medium (NGM) plates seeded with *Escherichia coli* OP50 and cultured at 20°C under standard laboratory conditions. Young adult animals were used for microinjection. The strains and alleles generated in this study along with the corresponding crRNA sequences are listed in Table 1.

#### B. Generation of ssDNA donor

Using either the dual PCR method or the restriction enzyme method, generate >8 µg of hemi-phosphorylated dsDNA (described in Figure 1). For the dual PCR method, we perform 500 µl PCR-II reaction, and purify using a single QIAquick PCR purification kit (QIAGEN: 28104), eluted in 30 µl (concentration usually in the range of 300–400 ng/µl).

#### C. λ-exonuclease treatment

Set up the following reaction:

| Component | Amount/Volume |
| --- | --- |
| 1. 10X λ-exonuclease buffer | 5.0 μl |
| 2. dsDNA template | > 8 μg |
| 3. λ-exonuclease (NEB: M0262) | 2.0 μl |
| 4. Nuclease-free H<sub>2</sub>O | to 50 μl |

Incubate the reaction at 37°C for 30 minutes and stop the reaction by using 0.5 M EDTA (2 µl). Purify ssDNA using spin columns and elute in 30 µl elution buffer.

*Note: In our hands, any purification kit with binding affinity of more than 5 µg effectively purified λ-exonuclease-digested DNA*.

*Note: Aim to elute ssDNA at a concentration greater than 100 ng/µl at the end of this step. We have tested injecting ssDNA in the range of 100 ng/µl to 250 ng/µl and not found a significant difference in editing efficiency*.

#### D. Preparing injection mix

1. Ribonucleoprotein (RNP) complex assembly Assemble the following components in this order prior to adding donor DNA:

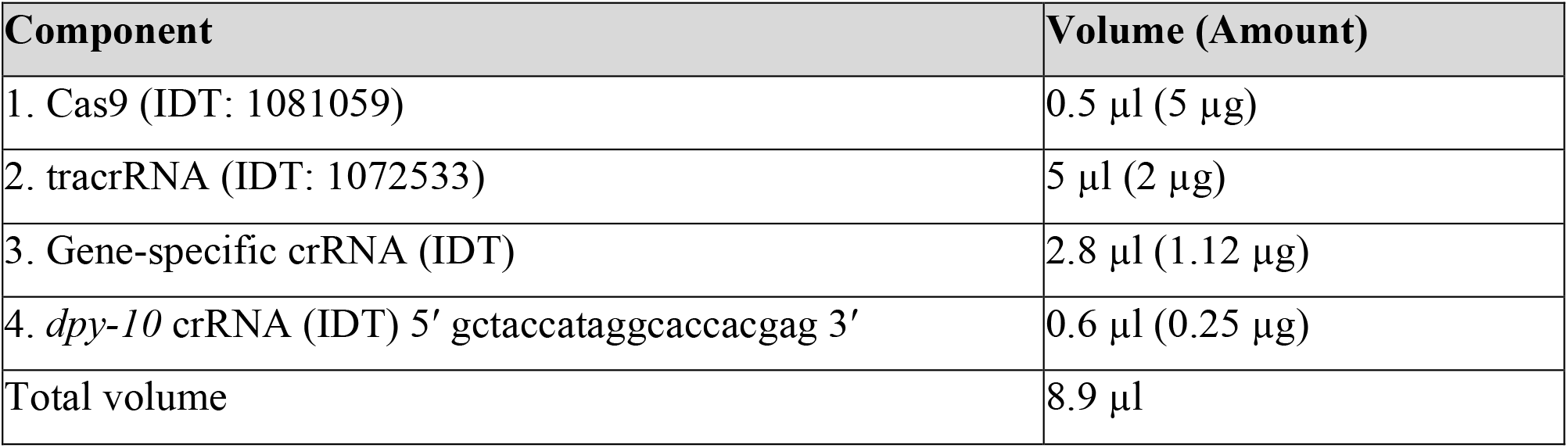 Incubate the mix at 37°C for 15 minutes to obtain the RNP complex.
2. During RNP incubation, take the ssDNA donor obtained from step C and prepare the DNA component mix:

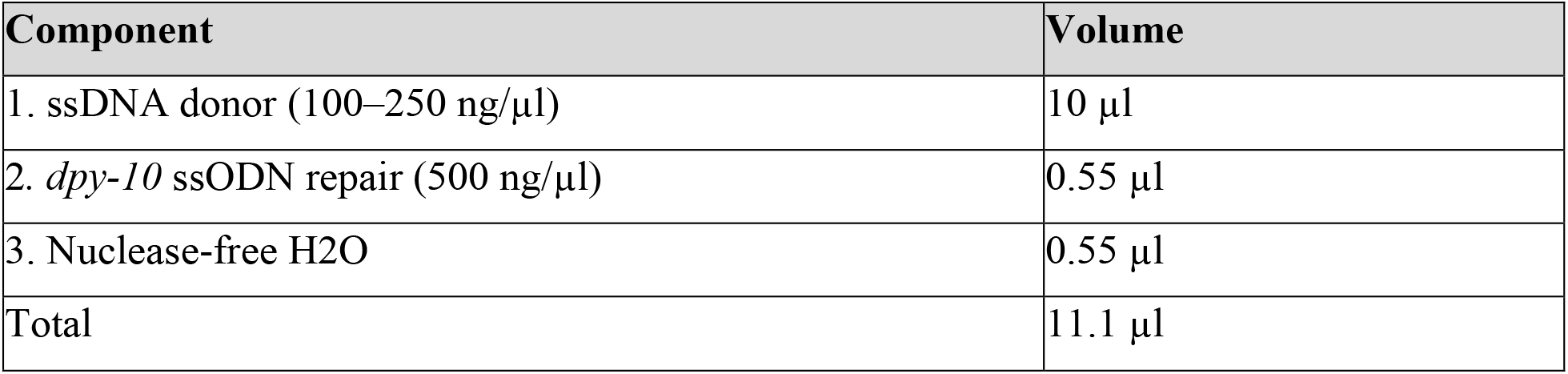 Subject the DNA component mix to a melting cycle in a thermal cycler as previously described (95°C – 2:00 min; 85°C – 10 sec, 75°C – 10 sec, 65°C – 10 sec, 55°C – 1:00 min, 45°C – 30 sec, 35°C – 10 sec, 25°C – 10 sec, 4°C – hold) [4].
3. Add the entire 11.1 µl of this melted product to the pre-incubated RNP assembly to bring the total injection mix volume to 20 µl.
4. Spin the injection mix at 15,000 rpm for at least 2 minutes to avoid needle clogging. Proceed to load needles.

#### E. Microinjection

Inject about 10 P0s (depending on the efficiency of the microinjector). Let the injected P0s grow for 3 days at 25°C.

#### F. Screening for edits

Identify the injected P0s that throw a high fraction of roller/dumpy animals [6]. Selecting from the top two to three plates, single about 30–50 roller/dumpy worms in total.

*Note: Prefer picking roller worms, as they are heterozygous for the dpy-10 mutation, which can be selected out in the next generation without the need for additional outcrossing* [6].

#### G. Genotyping of edited progeny

Screen for your desired edits from this cohort using fluorescent reporters (if possible), or by genotyping using PCR. For PCR-based genotyping, it is suggested to use one internal primer specific to the inserted cassette and the other primer specific to the gene locus that is outside of the homology arms used for the repair.

#### H. Fluorescence imaging

For fluorescence imaging, animals were mounted on agarose pads and immobilized using 5 mM Levamisole. Fluorescence images were acquired using a Nikon NiE fluorescent confocal microscope. Images were processed and adjusted for display using ImageJ Fiji software.

[utabl]

#### I. Data analysis

Following microinjection, progeny exhibiting the *dpy-10* co-conversion phenotype were selected for screening. Candidate animals were screened for the presence of the desired insertion by PCR or fluorescence, as appropriate for the construct. Editing efficiency was calculated as the number of animals carrying the confirmed insertion divided by the total number of *dpy* animals screened, expressed as a percentage.

Scatter plots were generated to visualize the relationship between editing efficiency and homology-arm length, insertion size, and other donor parameters. Pearson correlation analysis was used to assess the relationship between editing efficiency and 5′ homology-arm length, 3′ homology-arm length, and insertion size. Each construct was treated as a single observation, yielding n = 11 constructs for these analyses. Pearson correlation coefficients (r) and corresponding two-tailed p-values were calculated using n − 2 degrees of freedom. A p-value < 0.05 was considered statistically significant.

#### J. General notes and troubleshooting

1. These comparisons of editing efficiency encompassed different genomic loci and therefore cannot account for locus-specific effects on repair efficiency.
2. In case of limited success with editing at a particular locus, the crRNA-mediated dsDNA cut event can first be validated by genotyping PCR amplifying the region around the cut site. Up-shifts or down-shifts from the expected wild-type band would be suggestive of a positive cut event, but inefficient repair. Conversely, the absence of any shift in bands across F1 samples could suggest inefficiency of the crRNA.

### Validation of protocol

We show that this workflow can efficiently generate error-free knock-ins across multiple loci. For example, insertion of the split GFP-encoding sequence [11] in the *inx-5* gene locus (*inx-5(amz25[inx-5::splitGFP11X5])*) showed punctate INX-5::GFP localisation with pan-glially expressed, transgenic split-GFP(1–10) fragment, reminiscent of gap junction puncta (Figure 2C, D), validating the successful generation and expression of the intended knock-in.

With these new developments in the genome editing method, we asked whether reducing the homology-arm length could maintain high editing efficiency while simplifying donor preparation and reducing the associated primer synthesis costs and preparation time. We evaluated editing-efficiency across multiple genomic loci using λ-exonuclease-generated ssDNA donors with varying homology-arm lengths (Figure 1C). We examined the relationship between the length of the 5′ and 3′ homology arms and the editing efficiency. Across the constructs tested, editing efficiency did not show a significant correlation with 5′ homology-arm length (Pearson r = −0.12, p = 0.733) or 3′ homology-arm length (r = −0.21, p = 0.540) (Figure 2A). Although these comparisons encompassed different genomic loci and therefore cannot account for locus-specific effects, efficient editing was observed even with relatively short homology arms. Insertion of a ∼300 bp cassette encoding five tandem repeats of the 11^th^ beta-sheet of green fluorescence protein (*gfp*) [11] in the *inx-5* locus to generate the *inx-5(amz25[inx-5::splitGFP11X5])* allele was successfully achieved (editing efficiency of 47.9% among the screened *dpy* progeny) using 65 bp and 55 bp 5′ and 3′ homology arms, respectively. We also examined whether the size of the insert had any correlation with editing efficiency and observed no significant correlation across the constructs tested (Pearson r = −0.13, p = 0.709; Figure 2B). Together, these observations indicate that longer ssDNA homology regions are not necessarily associated with improved repair efficiency and suggest that homology regions exceeding 100 bp on each side are not an absolute requirement for successful repair using this approach. Our finding that efficient editing can be achieved with shorter homology regions facilitates simplification of the design and preparation of ssDNA donors.

Together, these results provide a simplified approach for generating ssDNA donors for CRISPR/Cas9-mediated knock-ins in *C. elegans*.

## Author contributions

A.V. and A.B.: Conceptualization, Writing; A.V., A.S., M.X.M., M.S., and Ananya Bandyopadhyay: Investigation; A.B.: Funding acquisition, Project administration, Supervision.

## Acknowledgments

We thank Selvanayaki Eswaramoorthy for expert assistance in *C. elegans* microinjections. NCBS *C. elegans* facility, Caenorhabditis Genetics Center (CGC), and wormbase.org for resources; members of the Bhattacharya lab for comments on this manuscript.

## Ethical considerations

All experiments used the nematode *Caenorhabditis elegans*, an invertebrate model organism that does not require institutional ethics committee approval for standard laboratory husbandry and microinjection procedures.

